# Flexible movement kernel estimation in habitat selection analyses with generalized additive models

**DOI:** 10.1101/2024.06.27.600970

**Authors:** Rafael Arce Guillen, Jennifer Pohle, Florian Jeltsch, Manuel Roeleke, Björn Reineking, Natasha Klappstein, Ulrike Schlägel

## Abstract

1. Habitat selection analysis includes resource selection analysis (RSA) and step selection analysis (SSA). These frameworks are used in order to understand space use of animals. Particularly, the SSA approach specifies the space availability of sequential locations through a movement kernel. This movement kernel is typically defined as the product of independent parametric distributions of step lengths (SLs) and turning angles (TAs). However, this assumption may not always be plausible for real data where short SLs are often correlated with large TAs and vice versa.
2. The objective of this paper is to relax the need for parametric distributions using *generalized additive models* (GAMs) and the R-package mgcv, based on the work of Klappstein et al. (2024). For this, we propose to specify the movement kernel as a bivariate tensor product, rather than independent distributions of SLs and TAs. In addition, we account for residual spatial autocorrelation in this GAM-approach.
3. Using simulations, we show that the tensor product approach accurately estimates the underlying movement kernel and that the fixed effects of the model are not biased. In particular, if the data are simulated with a copula distribution for SL and TA, i.e. if the independence assumption for SL and TA does not hold, the GAM approach produces better estimates than the classical approach. In addition, including a bivariate tensor product in the model leads to a better uncertainty estimation of the model parameters and a higher predictive quality of the model.
4. Incorporating a bivariate tensor product solves the problem of assuming parametric distributions and independence between SLs and TAs. This offers greater flexibility and makes the analysis of real data more reliable.

## 1 Introduction

Understanding the determinants of animal movement decisions using telemetry data is essential for conservation efforts and gaining insights into how animals respond to environmental resources. This analytical framework is commonly referred to as habitat selection analysis. The idea of this framework is to understand space use based on spatial features (Nathan et al. 2008, Forester et al. 2009, Kays et al. 2015, Northrup et al. 2022).

The primary aim of employing habitat selection models on telemetry data is to quantify the impacts of features influencing movement decisions, accounting for uncertainties, and considering the general movement capacity between sequential animal locations (space availability) (Northrup et al. 2022). Historically, telemetry data was collected at a relatively coarse temporal resolution, leading to the assumption of uniform availability between sequential locations across the entire study area. Consequently, treating observed animal locations as independent was deemed reasonable, and the prevalent approach involved the use of the Resource Selection Analysis (RSA) framework (Boyce et al. 2002, Manly et al. 2007). However, with technological advancements, locations can now be obtained at a finer time scale (Thurfjell et al. 2014), rendering the assumption of independence between sequential locations implausible. A detailed summary of animal movement models can be found at Hooten et al. (2017).

To address this dependency represented by the limited physical movement capacity of animals between short time intervals, Fortin et al. (2005) proposed a conceptual restriction of the availability domain at each time point based on observed step lengths (SLs; the Euclidean distance between sequential locations) and turning angles (TAs; the change in direction between sequential steps). The availability domain represents the spatial area that an animal is able/likely to physically reach from one point time point to the next one. Forester et al. (2009) expanded on this concept by incorporating SLs as linear or non-linear (smooth) effects in the model to correct for the fact that users do not know how animals would truly move in an homogeneous landscape. Frameworks accounting for the availability between sequential locations are known as Step Selection Analysis (SSA). Although the idea of including SLs as a non-linear smooth effect has not been widely adopted, Avgar et al. (2016) formalized mathematically the integration of SLs as linear effects. They assumed that availability is explained by a parametric movement kernel, defined as the product of an SL- and TA kernel, under the assumption of independence between SLs and TAs. For certain parametric distributions, the corresponding parameters can then be estimated as linear effects of SL and TA and transformations of them like log (SL) and cos (TA) (Avgar et al. 2016). With this, it became possible to include interactions between movement capacity modelled through SL/TA and spatial covariates. Conceptually, this methodology, termed Integrated Step Selection Analysis (iSSA), includes at each time point the assignment of a selection strength value for each location in the whole study area based on the last two locations. These values are used to calculate the likelihood to select that particular location according to the movement capacity of the animal, which is represented by the movement kernel. For instance, Roeleke et al. (2022) tracked 81 aerial-hawking bats in Uckermark, Germany, demonstrating evidence of their tendency to adjust their distance relative to the hunting or searching activity of conspecifics. Alston et al. (2020) examined the impact of temperature on the movement patterns of moose (Alces alces). Their findings suggested that increasing temperatures were associated with shorter movement patterns, a heightened preference for shaded areas, and negative selection of bog habitats.

Traditionally, iSSA is fitted using a conditional logistic regression (Forester et al. 2009, Avgar et al. 2016), which presupposes parametric exponential family distributions for SLs and TAs. This is typically a gamma distribution for SLs and a von Mises distribution for TAs. Although it has been shown to be beneficial for parameter estimation to account for movement (Forester et al. 2009), current iSSA approaches are limited for real data analysis in two ways: i) movement may not be adequately captured by parametric distributions, and ii) SLs and TAs are assumed to be independent, which might not represent animal behaviour in which usually there is dependence in movement variables (Morales et al. 2004). Hodel & Fieberg (2022) investigated the latter issue and its consequences, proposing a solution using the cylcop package (Hodel & Fieberg 2021) to simulate and model possible correlations between SLs and TAs with the aid of copulas.A copula is defined as a multivariate cumulative distribution function, which accounts for possible dependencies between the marginal distributions (Sklar 1973). In the SSA context, the movement kernel would thus be understood as a bivariate parametric distribution function of SLs and TAs. This adds complexity to the model and still relies on parametric assumptions, which are again not necessarily met for real data.

Besides the modelling of the movement kernel, assuming only linear effects of the spatial features may not always be suitable. The effects of these features could be better explained by a non-linear smooth function. For instance, it is not plausible that temperature has a linear effect on acceleration of hares since it is natural to expect less movement at extreme temperatures (Stiegler et al. 2023).

To address these issues, we suggest fitting habitat selection models using Generalized Additive Models (GAM) (Arce Guillen et al. 2023b, Klappstein et al. 2024). Following, the model implementation from Klappstein et al. (2024), we use the Cox proportional hazards likelihood. However, but additionally specifying the movement kernel as a two-dimensional smooth function *s*(SL, TA), which is a tensor product and which accounts for correlation between SLs and TAs. Thus, the main contribution of this paper is the inclusion of a tensor product and therefore, to represent different movement kernels and their possible correlation structures. For this, we use the mgcv R-package (Wood 2011). Similar to Arce Guillen et al. (2023) and Klappstein et al. (2024), we account for missing spatial variation to avoid the problems linked to spatial autocorrelation as explained by Arce Guillen et al. (2023).

By specifying SSA as a GAM-like model, users can specify the movement kernel without assuming any specific parametric distribution. Moreover, they can directly account for dependence between SLs and TAs via tensor products, and account for non-linear relationships between movement decisions and spatial covariates. Similar to the GF-iSSA (Gaussian Field iSSA; Arce Guillen et al. 2023), the implementation of spatial random effects can be included in the analysis (Klappstein et al. 2024). We demonstrate how to implement this and evaluate the performance of this flexible model in comparison to the classical iSSA in different scenarios. We also show (Supplemental Information) that, as a side benefit of the flexible GAM-formulation of the iSSA, the RSA may no longer be conceptually necessary for sequential movement data. If the movement kernel is flexible enough to represent a uniform distribution, users do not need to consider whether RSA or SSA should be applied to their tracks.

## 2 Materials and Methods

### 2.1 Model description

Conceptually, animal movement is driven by two main processes (Forester et al. 2009, Avgar et al. 2016, Hooten et al. 2017):

1. The movement kernel *ϕ*:

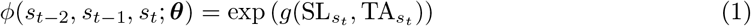
2. The habitat selection function (HSF):

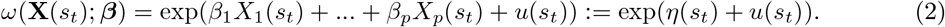

The movement kernel *ϕ* is used the represent the movement behaviour of animals in an homogeneous landscape and it depends on the time resolution of the data Forester et al. (2009). Here, 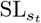 represents the SL of location *s*_*t*_ calculated in relation to the last observed location *s*_*t−*1_. Moreover, 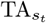 represents the TA at location computed based on the last two observed locations *s*_*t−*1_ and *s*_*t−*2_. In addition, ***θ*** represents potential parameters of the movement kernels, which could represent different statistical distributions. The function *g*() captures animal preferences for SLs and TAs jointly and is the function, which is later estimated with help of a bivariate tensor product. We name this model extension Tensor Product Integrated Step Selection Analysis (Tensor-iSSA).

The HSF *ω* describes the selection behaviour based on the spatial features Here, *β*_*j*_ denotes the selection strength regarding spatial feature *X*_*j*_(*s*_*t*_), and *u*(*s*_*t*_) represents the missing spatial variation, i.e. all spatial covariates, which were not available or collected by the user (Arce Guillen et al. 2023).

We adopt the same model formulation for the iSSA as Forester et al. (2009) and Avgar et al. (2016). The spatial density for the observation location *s*_*t*_ at time *t*, given the last two observed locations *s*_*t−*1_ and *s*_*t−*2_, is expressed as:

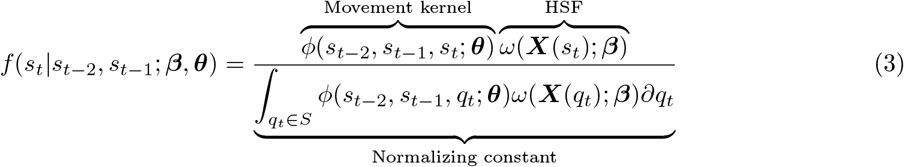

The joint log-likelihood for the full model is specified as the sum of conditional nonhomogeneous Poisson processes (NHPPs) (Arce Guillen et al. 2023), which simplifies to the cox Proportional hazards model used by Klappstein et al. (2024) after discretization (see Section 2.2 for details). The product of the movement kernel and the HSF is called Step Selection Function (SSF) (Forester et al. 2009, Avgar et al. 2016).

### 2.2 Model fitting

The integral of the step selection likelihood (Equation (3)) is intractable, and it is typically approximated via Monte Carlo integration (Michelot et al. 2024). That is, each observed location *s*_*t*_ is matched with *N* random points *{q*_1*t*_, *q*_2*t*_, …, *q*_*Nt*_*}* to derive an approximate step likelihood,

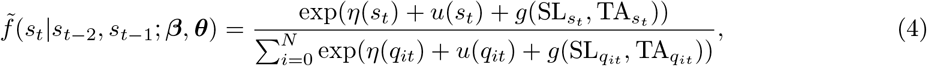

where *q*_0*t*_ is the observed location: *q*_0*t*_ = *s*_*t*_. We fit *u*() and *g*() as splines, which does not change the linearity of the model needed in GAM. The likelihood of the full movement path is the product of all step likelihoods, and this is equivalent to the likelihood of a Cox proportional hazards model, which can be implemented in mgcv (as explained in Klappstein et al. 2024).

Consistent with the methodology of Arce Guillen et al. (2023), we sample integration points uniformly within disks centred on each observed location, with a radius equal to at least the maximum observed SL. The use of nonuniform random integration points based on an *a priori* defined movement kernel, as usually used in iSSA, would lead to the estimation of a bivariate function for correction parameters rather than the original movement kernel. Only combined with the parameters of the assumed initial distribution, this would lead to movement kernel estimates, as highlighted by Avgar et al. (2016) and Klappstein et al. (2024). Therefore, we restrict the integration points to those sampled within disks (Fig. 1). Note that based on Equation 3, we could also sample at each time point integration points over the whole spatial domain. However, this would result in a significant loss of numerical speed. Furthermore, the likelihood contribution of spatial points further away than the maximum SL is negligibly small. Thus, our computational strategy not only saves time but also avoids introducing parameter bias (Arce Guillen et al. 2023, Michelot et al. 2024).

**Figure 1.**
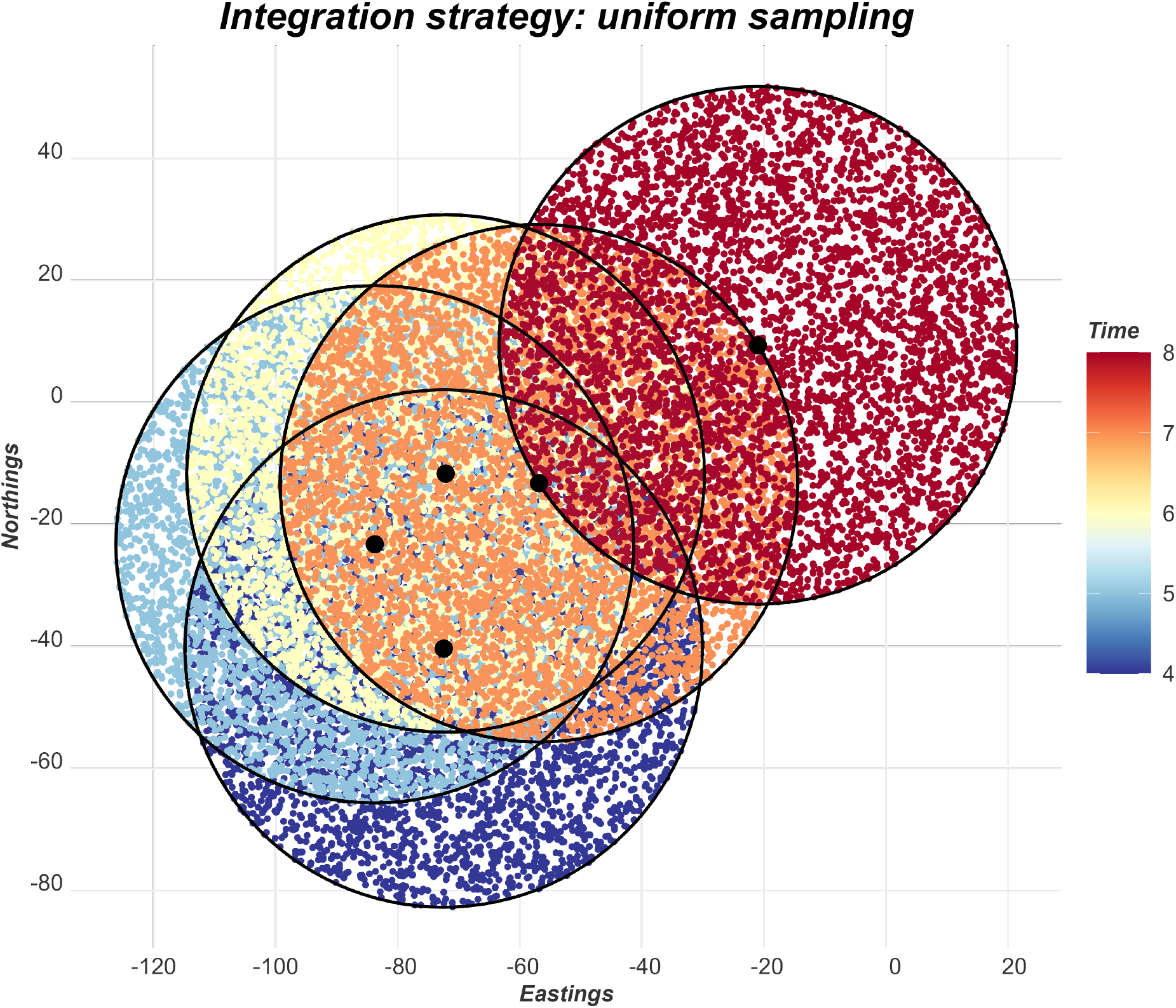
Tensor-iSSA: Integration strategy. At each time point, the domain of integration is restricted by a disk centred on the observed location (black dots). The radius of the disks is equal to at least the maximum observed SL. Integration points (colored dots) are then sampled uniformly over the disks of availability.

We fit the model using the GAM approach. GAMs are semi-parametric models that allow for estimating smooth functions using a pre-specified number of basis functions. The GAM approach is particularly useful when a non-linear relationship between a predictor and the dependent variable is expected. The movement kernel can naturally be interpreted as a non-linear bivariate function that encapsulates the correlation structure between SLs and TAs. Consequently, Generalized Additive Models (GAMs) serve as a convenient tool to model this intricate relationship. Furthermore, within this framework, non-linear spatial features can be effectively characterized by incorporating smooth functions. We employ penalized smooth terms to prevent overfitting (Wood 2011), executing our analyses with the mgcv package and the gam() function. Other functions within the mgcv package can also be employed to fit or extend this model.

We define the tensor product characterizing the movement kernel as a combination of a B-spline for the SLs (Eilers & Marx (1996), Wood (2017)) and a cyclic cubic regression spline for the TAs to account for the circular nature of TAs (Wood (2017)), accounting for their distinct units. Different penalization parameters are automatically estimated for SLs and TAs in the mgcv package. In addition, all the missing spatial features, represented by u(), are specified as a bivariate thin plate regression spline *s*(Longitude, Latitude).

### 2.3 Simulation study

We conducted simulations of animal tracks employing diverse movement kernel specifications. For this, we used a grid with a resolution 1001 *×* 1001 grid cells, and generated three landscapes. For each simulated step, we calculated the value of the step selection density based on the landscape features and corresponding movement kernel at all grid cells and then sampled one observed location correspondingly. For each configuration, we generated 100 animal tracks with 1000 observed steps. Table 1 provides an overview of the different simulation scenarios. All scenarios incorporate two continuous covariates, *x*_1_ and *x*_2_ (effect sizes equal to 1.5 and -1), simulated as Gaussian Fields with a variance parameter equal to 1 and spatial range parameters equal to 25 and 15 space units respectively. For this, we used the fields package (Nychka et al. 2021). Animals were assumed to have a centralizing tendency, modelled through a third spatial covariate *cen* defined as the distance to a center (0,0) with an effect size equal to -0.02. For simplicity and to isolate the marginal effect of the Tensor-iSSA, we did not introduce missing spatial variation in the simulation setting.

**Table 1.** Simulation setting. Description of the four simulated movement kernel scenarios and its parameters.

| Simulation setting | Description |
| --- | --- |
| Gamma(bimodal)-von Mises | SL: Bimodal distribution consisting of the combination of two gamma distributions ($\text{shape}_1 = 10$, $\text{scale}_1 = 1$, $\text{shape}_2 = 50$, $\text{scale}_2 = 1$) with weights equal to $\frac{1}{4}$ and $\frac{3}{4}$ respectively.<br>TA: von Mises distribution (mean = 0 , concentration = 1). |
| Copula | SL and TA are simulated using a bivariate copula distribution. We used a Weibull distribution (shape = 3, scale = 7) for the SLs and a wrapped Cauchy distribution (location = 0, scale = 0.3) for the TAs. |
| Gamma-von Mises | SL: Gamma distribution (shape = 10, rate = 1).<br>TA: Von Mises distribution (mean = 0 , concentration = 1). |
| Weibull-von Mises | SL: Weibull distribution (shape = 4, scale = 4).<br>TA: von Mises distribution (mean = 0 , concentration = 1). |

The first scenario involved a bimodal distribution for SLs, representing situations where animals exhibited different SLs in distinct states, such as searching and travel behaviors. The SL distribution had a peak at 10 and at 50 spatial units, respectively. The second scenario employed a movement kernel simulated with the cylcop package, using a circular-linear copula with quadratic sections. The correlation between SLs and TAs is evident in the circular-linear copula distribution plot (Fig. 2). Notably, when animals exhibited small SLs, they tended to display larger TAs, resembling foraging behavior. Conversely, with larger SLs, animals tended to move more straightforwardly, representing the travel mode (Hodel & Fieberg 2022). The third scenario, the Gamma/Von Mises setting, assumed independence between SLs and TAs and is the most commonly assumed scenario used in the iSSA framework. The fourth scenario employed a Weibull distribution for the SL and a von Mises distribution for the TA. We simulated this scenario to observe how our method performs when the true movement kernel is close but not identical to a Gamma/von Mises setting.

**Figure 2.**
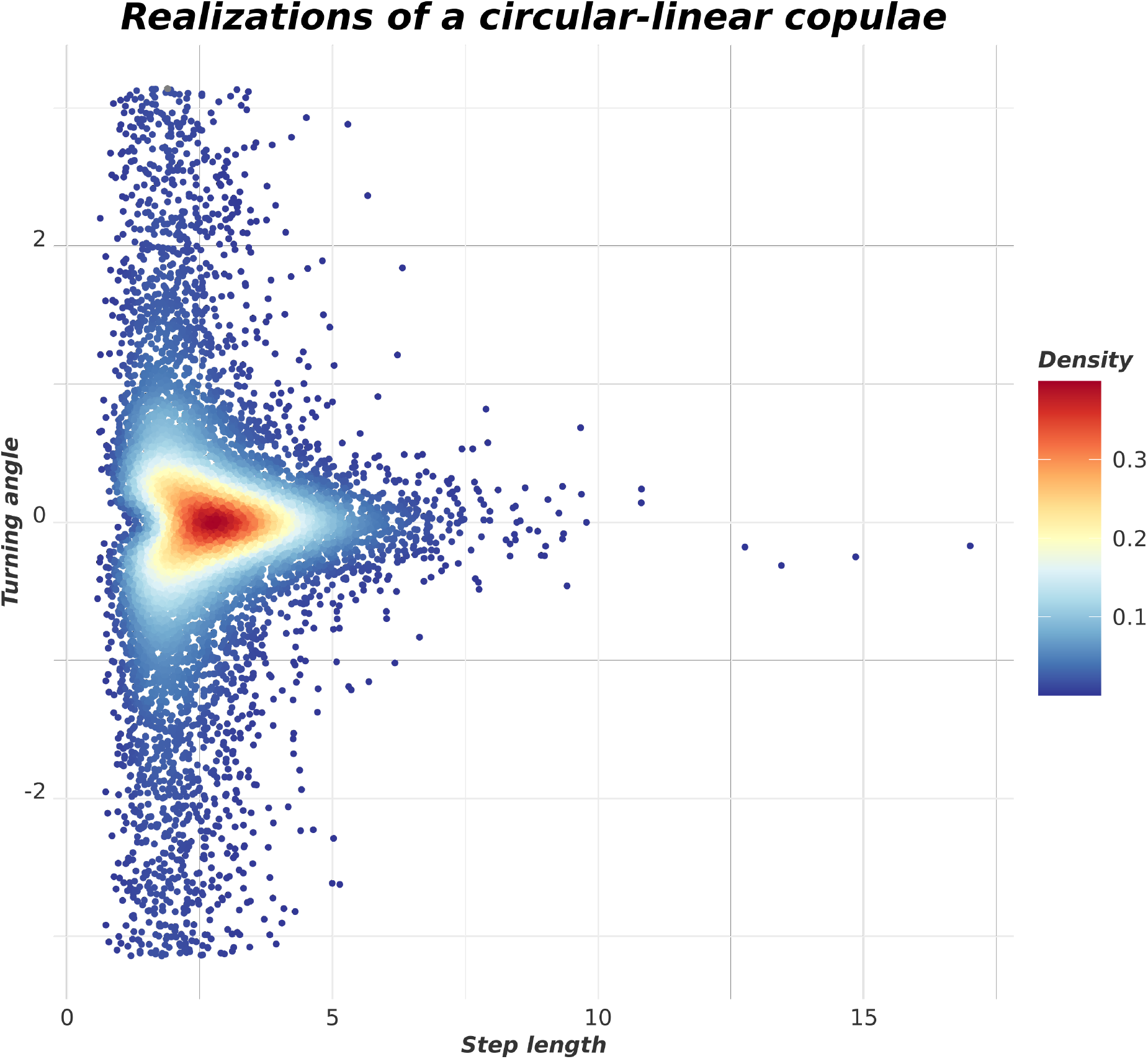
Second scenario: Density values of 10000 realizations of a Circular-linear copulae distribution with the cylcop package.

An SSF was fitted to each track using both standard iSSA (i.e., with gamma and von Mises distributions) and Tensor-iSSA approaches. We used 300 integration points, sampled uniformly on a disc for each observed location. For each scenario and model-fitting approach, we further fitted three model specifications: i) perfectly-specified models (i.e., with all covariates); ii) models without including the spatial variable *x*_2_ and iii) models without including *x*_2_ but having a spatial smoothing term accounting for missing spatial variation.

After fitting the model, we generated again 300 integration points within the disks of availability at each time point. After this, we calculated the true SSF values (the product of the true HSF and the true movement kernel) for these integration points and the predicted SSF values. After this, based on the true and predicted SSF values, we calculate the Mean Squared Error (MSE) to compare the predictive quality of the Tensor-iSSA compared to the iSSA method. In addition, we calculated the 95% statistical coverage of the spatial features parameters, i.e. the percentage for which the corresponding true values were covered by the 95% Wald confidence intervals respectively.

## 2.4 Case Study

We applied our method to 81 common noctule bats *Nyctalus noctula* tracked each for about 10 days in early or late summer of the years 2018 to 2020 in northern Germany (N 53.373945°, E 13.771231°) (Roeleke et al. 2022). The animal locations were sampled every eight seconds using the automated radio telemetry system ATLAS (Toledo et al. 2020). Flight paths for single individuals and nights were constructed from the raw data using filtering and smoothing approaches described in Roeleke et al. (2022). In this case study, we used our flexible tensor product approach adapted from Klappstein et al. (2024) to investigate habitat selection of bats during foraging flights. We used land use class as spatial covariate, which was obtained from a shapefile based on aerial infra-red photographs taken in 2009 (Land Brandenburg 2013). We specified the movement kernel as a tensor product and accounted for missing spatial covariates with bivariate thin plate regression splines (Wood 2017). Given the presence of outliers in the SLs, we used the corresponding 0.95-quantile of the observed SLs as the radius of the disks defining the domain of integration at each time point. The outliers were removed from the dataframe. We sampled the integration points uniformly, where each TA was sampled from U(*−π, π*) and each SL was the square root of a random draw from U(0, radius^2^) (Michelot et al. 2024). We sampled 100 integration points for each observed location and calculated the corresponding SLs and TAs with help of the amt package (Signer et al. 2019).

## 3 Results

### 3.1 Simulation study

In the Gamma(Bimodal)-von Mises scenario, we noted comparable estimates of the selection coefficients between methods, with small disparities observed between the complete (Including *x*1, *x*2 and *cen*) iSSA and the Tensor-iSSA approaches (Figure 5). In the copula scenario, however, it became evident that the selection coefficient estimates of the Tensor-iSSA were systematically less biased compared to the iSSA. This discrepancy arises because a simple Gamma/von Mises movement kernel does not accurately depict the complexity of the copula function, whereas the estimated movement kernel of the Tensor-iSSA aptly captures the correlation between SL and TA. In the Gamma-von Mises scenario, the iSSA accurately estimated the underlying process. Conversely, the Tensor-iSSA yielded selection coefficients nearly indistinguishable from those of the iSSA, indicative of the Tensor-iSSA’s ability for correctly modeling the movement kernel. This pattern extended to the Weibull-von Mises scenario. Despite being based on a Gamma-von Mises movement kernel assumption, the iSSA successfully approximated the true Weibull-von Mises movement kernel due to the notable resemblance between the gamma and Weibull distributions.

Introducing missing spatial variation by excluding *x*_2_ from the model specification induced a slight bias across all scenarios, albeit without a discernible trend. Incorporating this variation through a bivariate smoothing mechanism results in box-plots with more variability (Figure 3).

**Figure 3.**
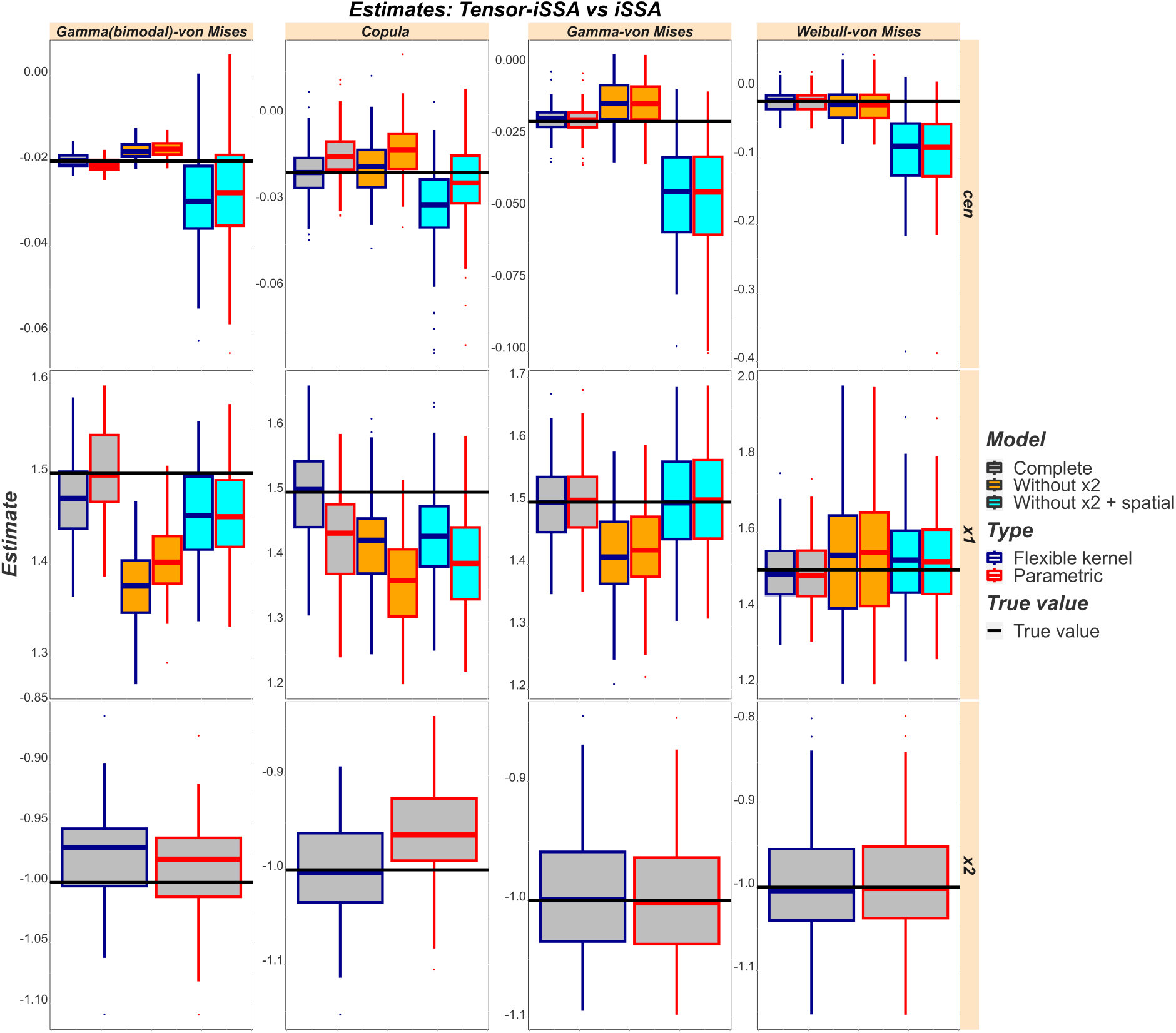
Selection strength results. The blue box-plots represent our tensor-product based approach (Flexible kernel). The red box-plots represent the classical iSSA approach (Parametric) with the Gamma/von Mises assumption.

Observing the summary of the selection coefficients derived from the fitted models along with their respective uncertainty estimates (Table 2), when missing spatial variation was introduced, thereby violating the independence assumption, we observed a low statistical coverage across all scenarios. This indicates an underestimation of the uncertainty associated with the selection coefficients by these models. Upon incorporating a bivariate smoothing technique to address the missing spatial variation, overall coverage improved in most cases, reaching more satisfactory levels. However, in the Gamma(bimodal)-von Mises, copula and Gamma-von Mises scenario, the coverage attained 100%, suggesting an overestimation of uncertainty. Furthermore, when considering the marginal effect of incorporating the tensor product, we observed that, in the copula case, the uncertainty estimation was superior in the Tensor-iSSA compared to the iSSA formulation. This improvement indicates that the tensor product mitigated residual autocorrelation resulting from the correlation between SLs and TAs.

**Table 2.** Selection coefficient estimates of the fixed effects. Each row summarizes 100 simulations. The quantiles are those of the estimates and do not represent the confidence intervals. The coverage represents the percentage for which the corresponding true values were covered by the corresponding Wald confidence intervals respectively.

| Scenario | Coefficient | Fit | Mean | True value | 0.05-quantile | 0.95-quantile | Coverage |
| --- | --- | --- | --- | --- | --- | --- | --- |
| **Gamma(bimodal)-von Mises** | cen | Tensor-iSSA | -0.02 | -0.02 | -0.02 | -0.02 | 0.97 |
|  | cen | Tensor-iSSA_x2 | -0.02 | -0.02 | -0.02 | -0.01 | 0.68 |
|  | cen | Tensor-iSSA_x2_spatial | -0.03 | -0.02 | -0.05 | -0.01 | 1.00 |
|  | cen | iSSA | -0.02 | -0.02 | -0.02 | -0.02 | 0.92 |
|  | cen | iSSA_x2 | -0.02 | -0.02 | -0.02 | -0.01 | 0.66 |
|  | cen | iSSA_x2_spatial | -0.03 | -0.02 | -0.05 | -0.01 | 1.00 |
|  | x1 | Tensor-iSSA | 1.47 | 1.50 | 1.40 | 1.54 | 0.93 |
|  | x1 | Tensor-iSSA_x2 | 1.38 | 1.50 | 1.31 | 1.44 | 0.22 |
|  | x1 | Tensor-iSSA_x2_spatial | 1.45 | 1.50 | 1.36 | 1.54 | 0.94 |
|  | x1 | iSSA | 1.50 | 1.50 | 1.44 | 1.57 | 0.98 |
|  | x1 | iSSA_x2 | 1.41 | 1.50 | 1.35 | 1.47 | 0.45 |
|  | x1 | iSSA_x2_spatial | 1.45 | 1.50 | 1.36 | 1.54 | 0.94 |
| **Copula** | cen | Tensor-iSSA | -0.02 | -0.02 | -0.03 | -0.01 | 0.94 |
|  | cen | Tensor-iSSA_x2 | -0.02 | -0.02 | -0.03 | -0.00 | 0.91 |
|  | cen | Tensor-iSSA_x2_spatial | -0.03 | -0.02 | -0.06 | -0.01 | 1.00 |
|  | cen | iSSA | -0.01 | -0.02 | -0.03 | -0.00 | 0.90 |
|  | cen | iSSA_x2 | -0.01 | -0.02 | -0.03 | 0.00 | 0.83 |
|  | cen | iSSA_x2_spatial | -0.02 | -0.02 | -0.05 | -0.00 | 1.00 |
|  | x1 | Tensor-iSSA | 1.50 | 1.50 | 1.39 | 1.63 | 0.96 |
|  | x1 | Tensor-iSSA_x2 | 1.42 | 1.50 | 1.30 | 1.53 | 0.84 |
|  | x1 | Tensor-iSSA_x2_spatial | 1.44 | 1.50 | 1.31 | 1.57 | 0.88 |
|  | x1 | iSSA | 1.43 | 1.50 | 1.32 | 1.54 | 0.81 |
|  | x1 | iSSA_x2 | 1.36 | 1.50 | 1.22 | 1.47 | 0.50 |
|  | x1 | iSSA_x2_spatial | 1.39 | 1.50 | 1.25 | 1.50 | 0.67 |
| **Gamma-von Mises** | cen | Tensor-iSSA | -0.02 | -0.02 | -0.03 | -0.01 | 0.95 |
|  | cen | Tensor-iSSA_x2 | -0.01 | -0.02 | -0.03 | -0.00 | 0.62 |
|  | cen | Tensor-iSSA_x2_spatial | -0.05 | -0.02 | -0.08 | -0.02 | 1.00 |
|  | cen | iSSA | -0.02 | -0.02 | -0.03 | -0.01 | 0.95 |
|  | cen | iSSA_x2 | -0.01 | -0.02 | -0.03 | -0.00 | 0.61 |
|  | cen | iSSA_x2_spatial | -0.05 | -0.02 | -0.08 | -0.02 | 1.00 |
|  | x1 | Tensor-iSSA | 1.50 | 1.50 | 1.39 | 1.59 | 0.96 |
|  | x1 | Tensor-iSSA_x2 | 1.42 | 1.50 | 1.30 | 1.54 | 0.66 |
|  | x1 | Tensor-iSSA_x2_spatial | 1.51 | 1.50 | 1.37 | 1.66 | 0.93 |
|  | x1 | iSSA | 1.50 | 1.50 | 1.40 | 1.60 | 0.97 |
|  | x1 | iSSA_x2 | 1.43 | 1.50 | 1.32 | 1.55 | 0.71 |
|  | x1 | iSSA_x2_spatial | 1.51 | 1.50 | 1.38 | 1.66 | 0.93 |
| **Weibull-von Mises** | cen | Tensor-iSSA | -0.02 | -0.02 | -0.04 | 0.00 | 0.94 |
|  | cen | Tensor-iSSA_x2 | -0.03 | -0.02 | -0.07 | 0.01 | 0.67 |
|  | cen | Tensor-iSSA_x2_spatial | -0.09 | -0.02 | -0.19 | -0.02 | 0.99 |
|  | cen | iSSA | -0.02 | -0.02 | -0.04 | 0.00 | 0.93 |
|  | cen | iSSA_x2 | -0.03 | -0.02 | -0.07 | 0.01 | 0.67 |
|  | cen | iSSA_x2_spatial | -0.09 | -0.02 | -0.19 | -0.02 | 0.99 |
|  | x1 | Tensor-iSSA | 1.49 | 1.50 | 1.35 | 1.62 | 0.96 |
|  | x1 | Tensor-iSSA_x2 | 1.53 | 1.50 | 1.29 | 1.79 | 0.68 |
|  | x1 | Tensor-iSSA_x2_spatial | 1.52 | 1.50 | 1.34 | 1.69 | 0.95 |
|  | x1 | iSSA | 1.49 | 1.50 | 1.34 | 1.63 | 0.95 |
|  | x1 | iSSA_x2 | 1.53 | 1.50 | 1.29 | 1.81 | 0.68 |
|  | x1 | iSSA_x2_spatial | 1.52 | 1.50 | 1.33 | 1.69 | 0.95 |

When assessing the predictive performance, the MSE scores were lower for the Tensor-iSSA than classical iSSA in all scenarios, except for the Gamma-von Mises scenario (Figure 4). However, even in such instances where the iSSA model perfectly conforms to a Gamma and von Mises distribution for SLs and TAs, our model exhibits almost identical MSE scores to the iSSA, indicating that Tensor-iSSA models may have generally better predictive performance over classical iSSA. In the copula and Gamma(bimodal)-von Mises scenarios, employing the movement kernel as a tensor product results in enhanced predictive performance compared to the iSSA approach when accounting for missing spatial variation. However, it’s crucial to note that such outcomes are contingent upon the specific simulation settings and should not be extrapolated as generalizations.

**Figure 4.**
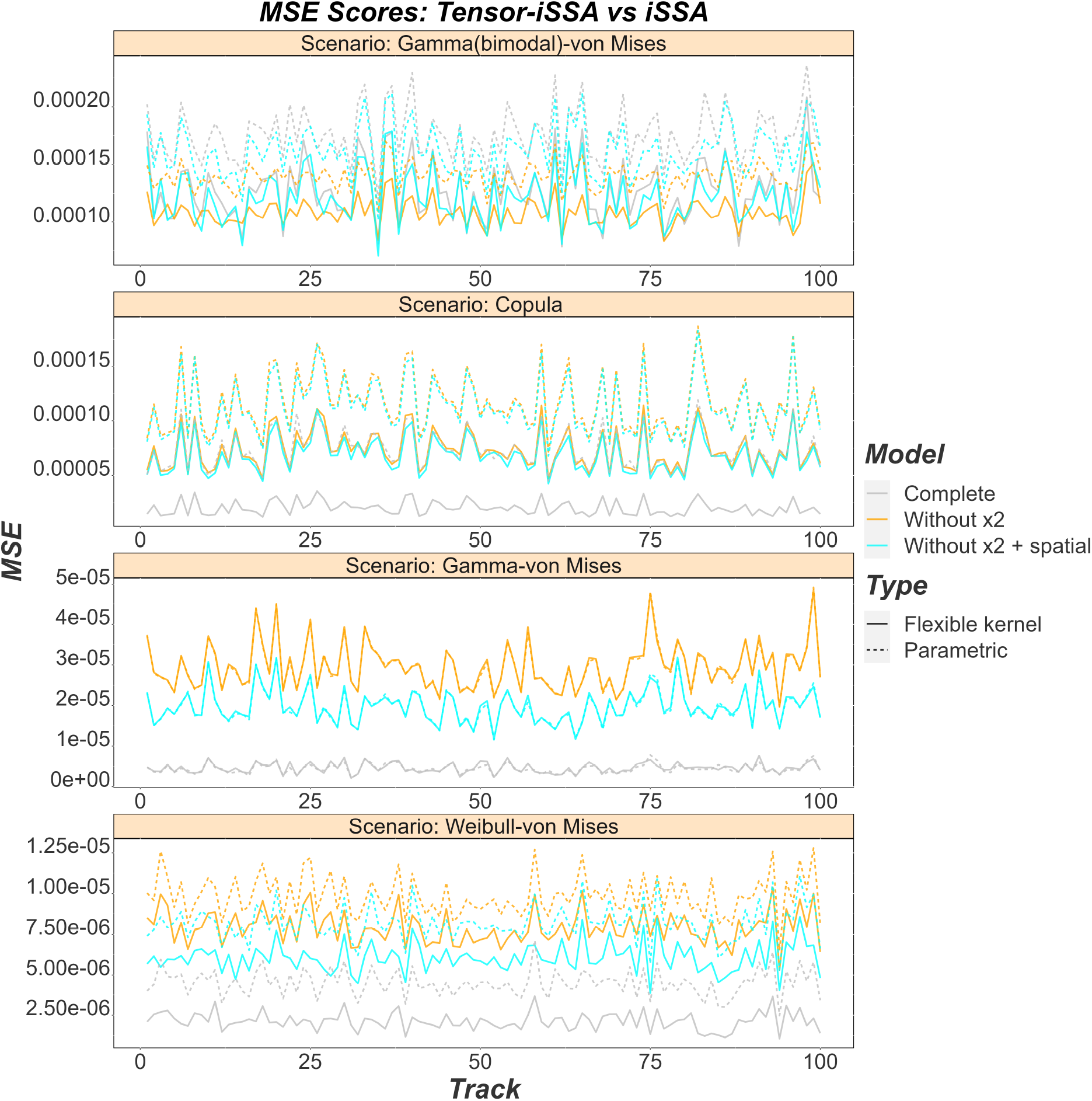
MSE scores for our different models. The continuous line represents the Tensor-iSSA model (flexible) while the dashed line represents the iSSA model (parametric). Lower values of the MSE indicate a better predictive quality of the SSF. The gray, yellow and blue colors represent the full model (complete), the model without *x*2 and the model without *x*2 but accounting for the missing spatial variation.

Finally, we examined the jointly estimated movement kernel across our four simulation settings (Figure 5). All simulated movement kernel structures were captured with Tensor-iSSA. It is evident that the two peaks characteristic of the Gamma(bimodal)-von Mises scenario were accurately captured, as well as the correlation between variables in the copula scenario. Similarly, the Gamma-von Mises and Weibull-von Mises settings exhibited the anticipated patterns (Figure 5).

**Figure 5.**
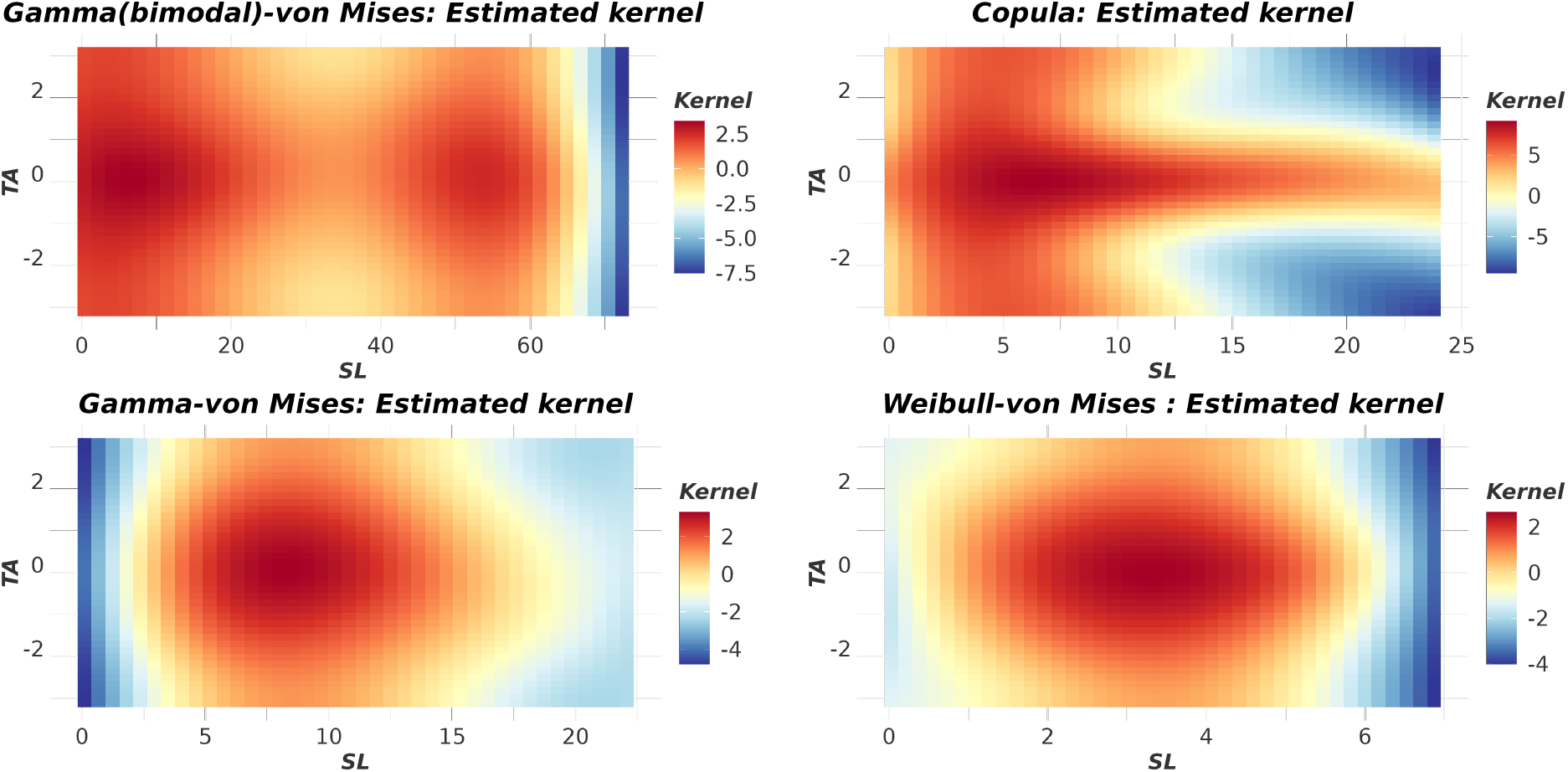
Tensor-iSSA (Flexible): Estimated movement kernel for the four different simulation settings derived from one simulated movement track each.

### 3.2 Case Study

Empirical data analysis revealed movement patterns similar to those found in copula distributions within the estimated movement kernel. Our model showed evidence that shorter steps of bats correlate with larger TAs, suggestive of exploratory behavior rather than straightforward navigation. In addition, longer SLs are typically associated with lower TAs, indicative of travel behavior (see Figure 6). However, even during travel phases, bats are likely to show occasional large directional shifts (i.e., TAs at |*π*|). Moreover, habitat selection coefficients showed that these bats select for standing water and swamps, compared to the reference category “agricultural land”(refer to Table 3 for full results). Furthermore, our model effectively captured the missing spatial variation, for which there is strong statistical evidence (see Figure 6). This model explained 31.8% of the deviance while the iSSA model with the Gamma-von Mises assumption explained 25.1% of it. Our model had also a lower AIC (1,579,911) compared to the iSSA model (1,644,252).

**Table 3.** Results of the Tensor-iSSA applied to bat data. Panel A. represents the parametric coefficients and panel B. represents the smooth therms. In this case the tensor product used to represent the movement kernel and the missing spatial variation represented by a bivariate smooth function.

| A. Parametric coefficients | Estimate | Std. Error | t-value | p-value |
| --- | --- | --- | --- | --- |
| Bush alley | -0.0484 | 0.0193 | -2.5124 | 0.0120 |
| Forest | -0.0513 | 0.0142 | -3.6139 | 0.0003 |
| Grassland | 0.0229 | 0.0123 | 1.8655 | 0.0621 |
| Greenland | 0.0075 | 0.0202 | 0.3697 | 0.7116 |
| Rural | -0.0669 | 0.0251 | -2.6649 | 0.0077 |
| Standing water_water | 0.1174 | 0.0140 | 8.3688 | < 0.0001 |
| Swamp | 0.1856 | 0.0224 | 8.2959 | < 0.0001 |
| Urban | -0.0153 | 0.0159 | -0.9616 | 0.3363 |
| B. Smooth terms | edf | Ref.df | F-value | p-value |
| te(sl_,ta_) | 17.1983 | 17.6986 | 158141.0442 | < 0.0001 |
| s(x2_,y2_) | 69.6102 | 85.5922 | 495.8596 | < 0.0001 |

**Figure 6.**
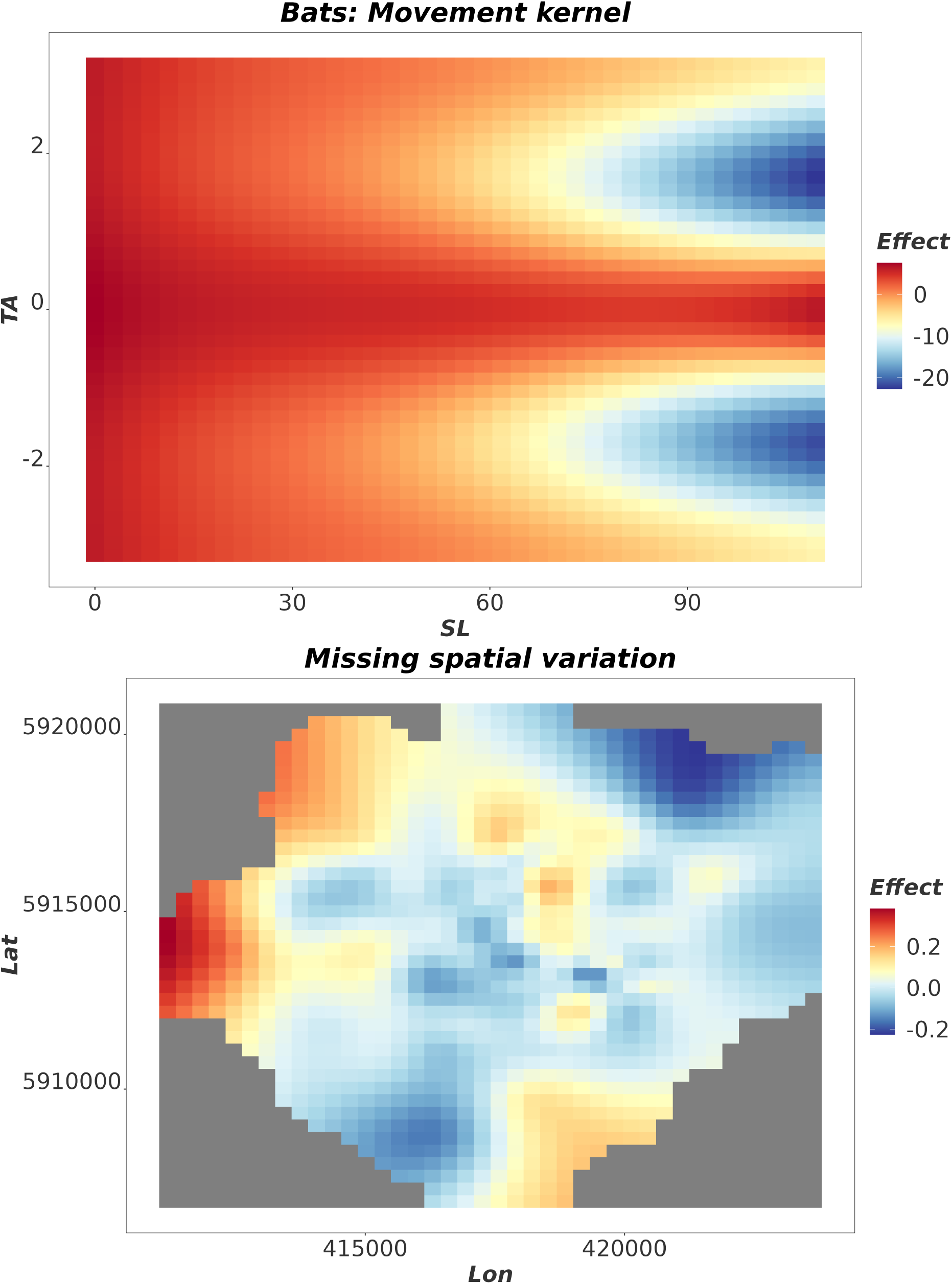
Common noctule bat case study results. The upper panel represents the estimated movement kernel (a tensor product of SL and TA) and the lower panel displays the missing spatial variation, estimated via a two-dimensional spatial smooth.

## 4 Discussion

In this study, we aim to alleviate the independence assumption between SLs and TAs of classical iSSA movement kernels. For this, we use the mgcv model implementation of Klappstein et al. (2024) and include a tensor product in the model to represent the movement kernel. Further, our approach does not rely on parametric distributions, leading to a straightforward and user-friendly implementation with the mgcv package (as in Klappstein et al. 2024). Using simulations and a real data example, similar to Hodel & Fieberg (2022), we show that the movement kernel may not always be adequately described by the product of independent Gamma and von Mises distributions. Instead, it manifests as a multifaceted bivariate function intertwined with various factors including time resolution, species characteristics, and environmental variables. The intricate interplay between SLs and TAs, characterized by their complexity and interdependence, finds an appropriate representation through the flexibility afforded by a tensor product. The Tensor-iSSA offers three main advantages. First, for scenarios, where SLs and TAs are correlated to each other (Copula), our approach leads to a bias reduction of the spatial covariates effects. Second, for all scenarios, the Tensor-iSSA offers a higher predictions quality of the SSF. Thus, users interested in predictions would benefit from this method. Third, in the copula case, our method leads to a better statistical coverage of the spatial effects suggesting a more accurate uncertainty estimation. In addition, similar to Arce Guillen et al. (2023), Klappstein et al. (2024), we offer the possibility to account for missing spatial covariates. This also leads to a better uncertainty estimation of the model parameters.

Our findings indicate that even when the data is simulated according to the Gamma-Von Mises framework, our model consistently provides accurate inference regarding the model parameters. Notably, it yields unbiased estimates of the selection coefficients, closely resembling those obtained from classical iSSA model, which is a correctly-specified model in this case. In contrast, while the iSSA model exhibits a slight bias in estimating the selection coefficients for the copula case, our model returns unbiased estimates. The remaining two scenarios exhibit similar patterns. However, upon evaluating the predictive performance of our model, we observe that it is at least comparable to the iSSA approach. In the Gamma-Von Mises case, both models demonstrate nearly identical MSE scores. Nevertheless, for scenarios involving Gamma(bimodal)-von Mises, copula, and Weibull-Von Mises distributions, our model exhibits superior predictive quality. Consequently, for users prioritizing predictive accuracy, we recommend employing the Tensor-iSSA. Finally, we discover that incorporating a bivariate smooth to account for missing spatial variation may, in certain cases, lead to an overestimation of the uncertainty associated with the model parameters. One potential remedy could involve modeling the missing spatial variation using a formal multivariate spatial Gaussian process, for example using the the approach proposed by Arce Guillen et al. (2023), instead of using a bivariate smooth. While feasible with the mgcv package, this lies beyond the scope of our present study. Note that for a more flexible modelling of the movement kernel of the iSSA approach, users could include also terms of SLs, TAs and interactions thereof in the model. However, they still rely on parameter assumptions.

The Tensor-iSSA revealed intriguing patterns in the case study. It estimated a movement kernel for bats that represented a similar structure to a copula distribution, displaying a clear correlation between SLs and TAs. Notably, the model indicated a peculiar behavior when bats were in travel mode, characterized by occasional large directional shifts instead of slight turns. This phenomenon may be attributed to the transition from foraging mode to traveling mode. This kind of estimation would not have been possible without a tensor product.

Users who favor the Poisson formulation of the iSSA model (Muff et al. (2020), Arce Guillen et al. (2023)) can also implement this model utilizing the mgcv package and the function bam() (Arce Guillen et al. 2023b). This functionality allows users the flexibility to execute the model using multiple nodes. However, it’s important to note that the inclusion of time-dependent intercepts may result in a reduction in numerical speed. As a result, this approach is primarily recommended for exceedingly complex models applied to very large datasets, where computational resources such as clusters are available for efficient fitting.

As highlighted by Arce Guillen et al. (2023), spatial models may be susceptible to spatial confounding, irrespective of whether terms are included to address missing spatial variation (Dupont et al. 2020, Thaden & Kneib 2018). Consequently, both models with and without the incorporation of a bivariate spline could potentially encounter this phenomenon. Nevertheless, Dupont et al. (2020) demonstrated a method for mitigating spatial confounding when the missing spatial variation is modeled as a bivariate smooth function, utilizing the capabilities of the mgcv package. Thus, when using spatial effects as in telemetry data and getting unexpected results, it might be worth to apply spatial confounding methods.

Our proposed approach models the correlation between SLs and TAs, which realistically captures behavioural dynamics of animals in conjunction with habitat selection (Morales et al. 2004). Besides a flexible modelling of the movement kernel and the inclusion of missing spatial variation, the Tensor-iSSA can be tailored to accommodate any number of smooth and random effects that can be specified in mgcv (as illustrated by Klappstein et al. 2024). For example, individual-level random effects can be used to account for inter-individual variability, and varying-coefficient models can be used to assess non-linear interactions. Additionally, Generalized Additive Mixed Models (GAMMs) with the gamm() function offer a versatile framework for specifying random slopes, thereby extending the modeling capabilities in capturing nuanced spatial dynamics.

In summary, the Tensor-iSSA leads to better predictions. It also can lead to a spatial effects parameters bias reduction and a better uncertainty estimation. In addition, it provides a user-friendly option to account for missing spatial variation, leading to a better uncertainty estimation. Users also benefit from the added flexibility of representing different movement kernel and spatial covariates with tensor products and smoothing functions respectively. Finally, the Tensor-iSSA can be used as an exploratory tool to gain insight into the complexity of the underlying structures.

## AUTHOR CONTRIBUTIONS

Rafael Arce Guillen, Jennifer Pohle, Ulrike E. Schlägel, Natasha Klappstein and Björn Reineking contributed conceptually and formulated the model. Rafael Arce Guillen and Natasha Klappstein provided the practical implementation of the model. Manuel Roeleke and Florian Jeltsch contributed with data and ecological concepts of the manuscript respectively. All authors assisted in writing and editing the manuscript

## ACKNOWLEDGEMENTS

This project was founded by the *German Research Foundation* (DFG) Grant SCHL 2259/1-1 granted to US and the BioMove Research Training Group (DFG-GRK 2118) lead by FJ.

## AUTHOR APPROVALS

All authors have seen and approved the manuscript. Moreover, this manuscript has not yet been accepted or published elsewhere.

